# Asymmetric stimulus representations bias visual perceptual learning

**DOI:** 10.1101/2023.07.11.548603

**Authors:** Pooya Laamerad, Asmara Awada, Christopher C. Pack, Shahab Bakhtiari

## Abstract

The primate visual cortex contains various regions that exhibit specialization for different stimulus properties, such as motion, shape, and color. Within each region there is often further specialization, such that particular stimulus features, such as horizontal and vertical orientations, are overrepresented. These asymmetries are associated with well-known perceptual biases, but little is known about how they influence visual learning. Most theories would predict that learning is optimal, in the sense that it is unaffected by these asymmetries. But other approaches to learning would result in specific patterns of perceptual biases. To distinguish between these possibilities, we trained human observers to discriminate between expanding and contracting motion patterns, which have a highly asymmetrical representation in visual cortex. Observers exhibited biased percepts of these stimuli, and these biases were affected by training in ways that were often suboptimal. We simulated different neural network models and found that a learning rule that involved only adjustments to decision criteria, rather than connection weights, could account for our data. These results suggest that cortical asymmetries influence visual perception and that human observers often rely on suboptimal strategies for learning.

## Introduction

The ability to discriminate between different visual stimuli is thought to depend on their visual cortical representations: Discrimination is easiest for stimuli that yield very different response patterns in neuronal populations (Figure 1A; Ahissar & Hochstein, 2004; Panzeri, Harvey, Piasini, Latham, & Fellin, 2017). Training-induced improvements in perceptual abilities, known as visual perceptual learning (VPL), have been suggested to arise from adjustments in sensory neuron tuning (the retuning hypothesis; (Karni & Sagi, 1991; Wenliang & Seitz, 2018)) or adjustment in the readout weights of the sensory neurons (the reweighting hypothesis; (Lu & Dosher, 2022; Sotiropoulos, Seitz, & Series, 2011)). Both theories assume *optimality* in VPL, which is to say that they involve learning that maximizes discrimination performance for a trained task.

**Figure 1.**
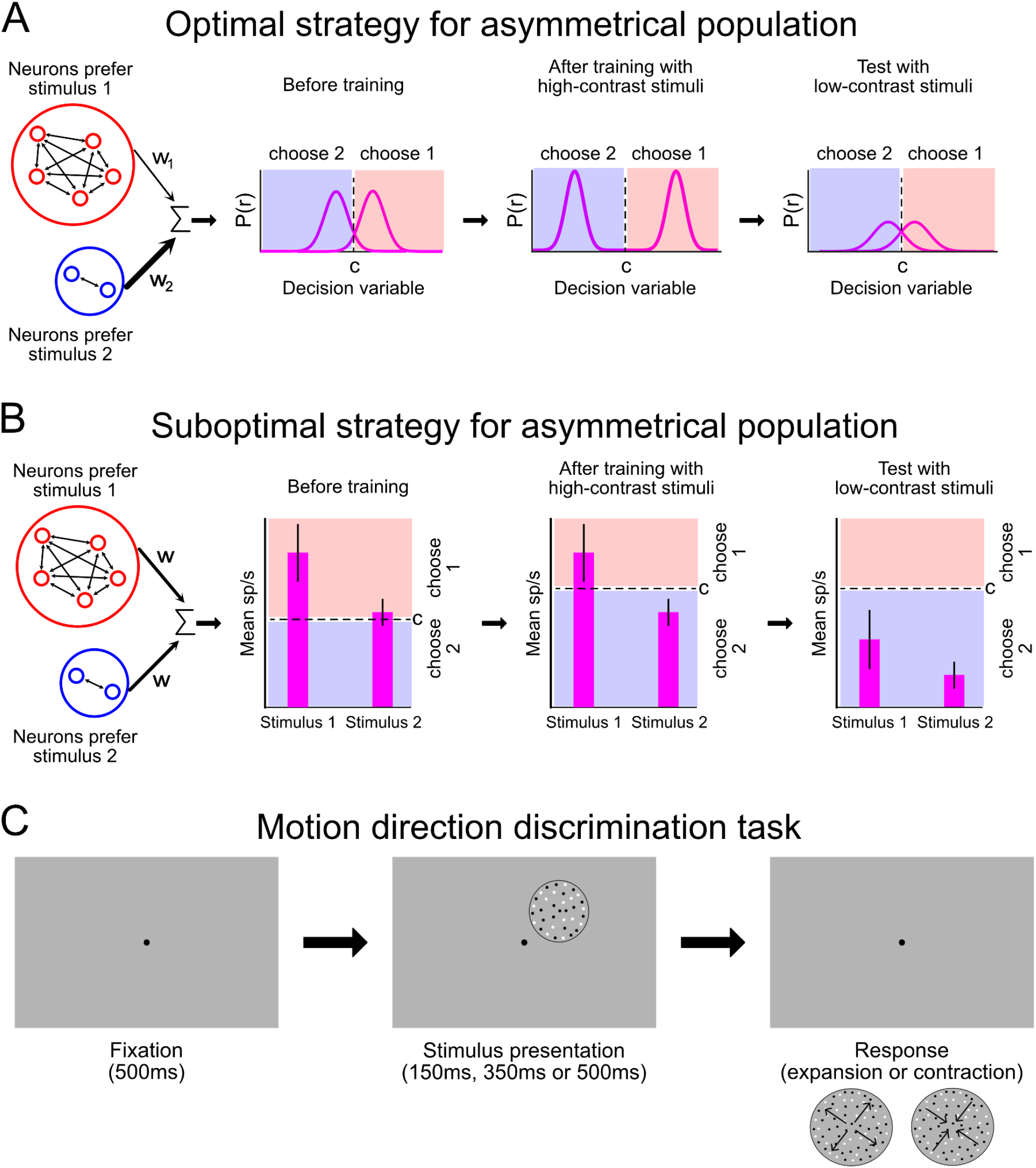
Schematics for the alternative readout strategies and the experimental procedure. A) The schematics of perceptual decision making using an optimal strategy. In an asymmetrical population (left), more neurons are selective for one stimulus condition (red) than for the other (blue). The population consists of neurons with a variety of sensitives and tuning properties to the two alternatives. Readout of the neuronal responses involves selecting relevant sensory signals that contribute to making a decision by weighting neurons based on their selectivity and summing the weighted outputs to compute the decision variable. This strategy can compensate for the asymmetry during readout, by assigning higher weights to stimulus with weaker representation or by selecting equal numbers of neurons from each pool. Training can alter the responses or the readout weights (center), thus preventing developing biased perceptual decisions (right). B) An alternate strategy is to sum the total population response, with equal readout weights for all neurons, yielding a scalar measure of the likelihood of one stimulus being present (left). During training, observers can adjust their decision criterion according to the properties of the stimuli (center), but this approach will yield consistent perceptual biases when the stimulus strength changes (right). C) Motion direction discrimination task used in the pre-training (one session), training (nine sessions) and post-training (one session) phases of the experiment. Each trial began when a fixation target appeared for 500 ms. Depending on the task’s difficulty, the stimulus appeared for 150 ms, 300 ms or 500 ms. The stimulus was contracting or expanding optic flow motion. Two options appeared on the screen when the stimulus and fixation target disappeared. The participant could report the direction of the motion (expansion vs. contraction) with the keyboard.

In certain perceptual situations, suboptimal learning strategies could also lead to perceptual improvements. Specifically, if the population response is *asymmetrical*, so that one stimulus yields higher neuronal responses than another (Figure 1B), learning can proceed by adjustment of decision criterion, without the need for retuning or reweighting of sensory responses. Such asymmetrical population responses are very common in visual cortex. Examples include stronger responses for cardinal orientations (Levick & Thibos, 1982; Nikara, Bishop, & Pettigrew, 1968; Schall, Vitek, & Leventhal, 1986; Vidyasagar & Urbas, 1982), for centrifugal motion stimuli (Albright, 1989), for horizontal disparities (DeAngelis & Uka, 2003), and for circular shapes (Dumoulin & Hess, 2007). A particularly clear example of this kind of asymmetry is the overrepresentation of expansion optic flow compared to other optic flow patterns in the medial superior temporal (MST) area of visual cortex (Duffy & Wurtz, 1995; Graziano, Andersen, & Snowden, 1994; Mineault, Khawaja, Butts, & Pack, 2012; Saito, Yukie, Tanaka, Hikosaka, Fukada, & Iwai, 1986).

Observers can change decision criteria easily when it is warranted by the task (Aberg & Herzog, 2012). However, this is a suboptimal strategy under most conditions because it is vulnerable to producing biased perceptual decisions when the stimulus strength deviates from that of the trained condition (Figure 1B). For example, training with a high-contrast stimulus might cause observers to increase their decision criteria, leading to perceptual biases for low-contrast stimuli. We therefore wondered whether observers would rely on suboptimal strategies when training with stimuli that have asymmetric cortical representations.

To answer this question, we trained a group of human observers on an optic flow two-alternative forced choice (2AFC) task that involved distinguishing between expansion and contraction optic flow. We found that observers exhibited biases that were predictable from the characteristics of the asymmetrical representation in MST and that these biases evolved with training in a way that was consistent with a suboptimal learning strategy.

To understand these dynamics, we compared neural networks that were trained to perform the same psychophysical task with different learning rules. Among them, only a suboptimal learning rule involving adjustment of decision criteria was able to account for the learning sensitivity and biases observed in humans both before and after training. These results suggest that asymmetric representations of the kind that are commonly found in the primate cortex can predict the properties of visual learning.

## Methods

### Observers and Apparatus

Fifty-six observers with normal or corrected-to-normal vision participated in this study (20 male observers, 36 female observers; age: 23.2 ± 3.36 years, range: 18-32 years). All observers were naïve to the purpose of the study and to visual psychophysics. Observers gave written, informed consent before their participation, and the study was approved by the Ethics Committee of the Montreal Neurological Institute and Hospital (NEU-06-033).

The experiment was run remotely and was controlled by a browser-based program (Article 19 Group; Montreal) that displayed the stimuli, monitored performance, and stored the data. Participants completed the study at home, on personal computers. Prior to the start of the experiment, participants were asked to provide the experimenter with their screen size. All visual displays were in the range of 13-27” in diagonal and had a refresh rate of at least 30 Hz. Stimulus parameters were calibrated to each observer’s screen size.

### Experimental procedure

The stimulus used in each experiment was an optic flow stimulus composed of an expanding or contracting random dot kinematogram (RDK). The stimulus was presented on a gray background in the upper right quadrant at an eccentricity of 6 degrees. The RDK was composed of small (0.06 degrees) black dots in a 3-degree radius aperture with a dot density of 2.6 dots/deg^2^. Dot velocity was set to 20°/sec. Stimulus duration and dot lifetime (after which the trajectory of the dot ended and was restarted at a random position) varied depending on the task’s difficulty. The dots presented were either “signal dots” or “noise dots”. Signal dots moved coherently in a specific direction, whereas noise dots moved in random directions. The coherence of the RDK stimulus refers to the proportion of signal dots.

Each trial started with a fixation point that the observer had to fixate for 500 ms. After the stimulus presentation (duration variable depending on the experimental condition), the fixation point and stimulus disappeared, and two options appeared on the screen (expansion and contraction). The observer was required to input their response with the keyboard (left for expansion, right for contraction). A green flash signaled a correct response and a red flash signaled an incorrect response. The direction of motion of the stimulus was chosen randomly for each trial.

As noted in the Introduction, we were interested in studying learning for different levels of task difficulty. Given the remote nature of the experiments, we were unable to precisely control stimulus contrast, which in any case does not matter much for optic flow tasks (Morrone, Burr, & Vaina, 1995). We therefore used stimulus duration to modulate task difficulty. For condition 1 (20 observers), the stimulus was shown for 150 ms with a dot lifetime of 75 ms. For condition 2 (19 observers), the stimulus was shown for 350 ms with a dot lifetime of 250 ms. For condition 3 (17 observers), the stimulus was shown for 500 ms with a dot lifetime of 250 ms. Each condition consisted of three phases: a pre-training phase, a training phase and a post-training phase. In every phase of the experiment, the observers completed a direction discrimination task in which they reported the direction of the motion of the stimulus (contraction or expansion) with the keyboard (Fig. 1B). Dot coherence was adjusted differently in each phase, as described below.

## Experimental phases

### Pre-training

The pre-training session required observers to report the direction of the motion of the stimulus at different coherence levels. The coherence levels tested were 0.025, 0.05, 0.1, 0.15, 0.2, 0.5 and 0.9. The pre-training session was composed of 490 trials. Each block of 70 trials tested the same coherence level. The order of coherences tested varied randomly for each participant. The pre-training session lasted approximately 30 minutes.

### Training

The training phase ran over nine days and required one training session to be completed per day. One training session was composed of 4 blocks of training. Each block was composed of 125 trials. The observers were compensated with 1.2 cents (Canadian) per correct response. At the start of each block, the initial coherence of the stimulus was set to 0.7. The coherence for each subsequent trial was set using a 2-down-1-up adaptive staircase procedure, resulting in an 83% convergence level (Leek, 2001). Observers were allowed to take a break between each block. Each daily training session lasted approximately 30 minutes.

### Post-training

The procedures for the post-training were the same as for the pre-training. The only difference was that observers received no feedback for correct and incorrect responses, in order to avoid further training effects. The order of coherences tested was chosen randomly and differed from the order of coherences tested in the pre-training phase.

## Data analysis

### Psychometric curve fitting

The observers’ performance as a function of coherence was characterized by fitting a Weibull function to the proportion of correct responses using the maximal likelihood algorithm (MATLAB Palamedes toolbox for analyzing psychophysical data, (Prins & Kingdom, 2018)). The Weibull function is given as:

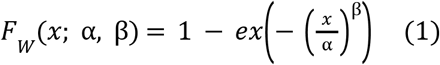

where αdetermines the threshold and parameter β corresponds to the slope of the function. The criterion of maximum likelihood was used to find the best-fitting psychometric function to each observer’s performance.

### Bias calculation

We used the equation below to calculate the bias (Wickens, 2001):

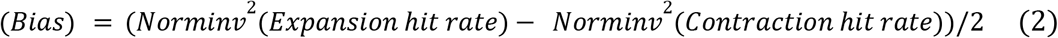

where Norminv is the Normal inverse cumulative distribution function. A log-transform was used to remove a nonlinear effect of bias. Negative bias values indicate a bias toward contraction, and positive values indicate a bias toward expansion.

### Statistical comparisons

Statistical comparisons of computed bias values were based on the one-tailed Wilcoxon signed-rank Test (WSR). In order to calculate the statistical difference of biases before and after the training, we used the Wilcoxon rank sum test (WRS).

### Model

To determine how observer learning might have progressed, we simulated 4 computational models, each starting from the assumption that psychophysical decisions were based on the asymmetrical representation found in area MST. The three models embodied different optimal and suboptimal learning rules. The equation below describes the model:

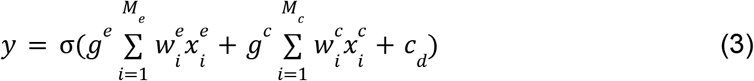

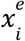 and 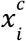 are the activation of the expansion and contraction neurons in MST, respectively. 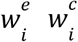 are their readout weights (the strength of synaptic connections to the readout neuron),*g*^*e*^ an *g*^*c*^ are sensory gains, *c*_*d*_ is the decision criterion (or the bias term), and *y* represents the activation of the readout neuron. In different versions of our models, the tuning properties of the neurons are fixed, and the readout weights 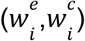, the decision criterion (*c* _*d*_), and the sensory gains (*g*^*e*,^*g*^*c*^) are the only model parameters that change. All the trainable parameters of the model (θ) were optimized by using the gradient descent learning rule:

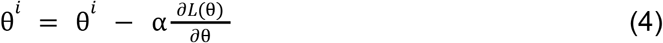

*L* is the error rate of the model which we quantified using a binary cross-entropy loss:

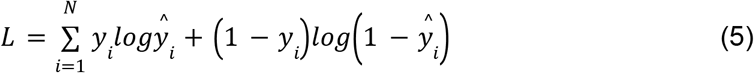

*ŷ*_*i*_ s and *y*_*i*_ s are the model outputs and the ground truth labels, respectively for the *i* trial.

All the synaptic weights 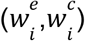 were initialized to one, assuming that at initialization, there is no a priori assumption about the relative importance of the sensory neurons in the task. The decision criterion *c*_*d*_ was initialized to zero, and the sensory gains (*g*^*e*^ *g*^*c*^) were initialized to one.

The output of each sensory neuron 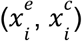 was modeled as a sigmoid function:

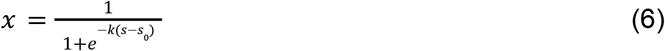

For the expansion and contraction neurons, the parameter *k* was set to model their opposite input selectivities (*k* = ±5, *S*_*0*_ =− 0. 5). Variability across sensory neurons was modeled by adding random gaussian noise to *k* and *S*_*0*_(ε∼*N*(0, 0. 2)).

## Results

### Human experiment

We trained 56 human observers to distinguish between contracting and expanding optic flow motion over the course of nine sessions and measured their perceptual bias before and after the training. To distinguish between different learning and readout strategies (optimal vs suboptimal), we divided the observers into three groups, each with a different stimulus duration (150, 350, and 500 ms). Stimulus duration modulated task difficulty, as shown below. Observers from all three conditions were tested before training (pre-training test) and after training (post-training test) to examine how perceptual learning changed their perceptual sensitivity and bias.

In all three duration conditions, training improved discrimination accuracy, as measured from a comparison between the mean performance on the pre- and post-training sessions (Fig. 2A). Across observers, the threshold for accurate performance (82%) was significantly reduced after the training in all three conditions (p = 0.02 for 150ms task, p < 0.001 for 350ms task and p = for the 500ms task, t-test; Fig. 2B).

**Figure 2.**
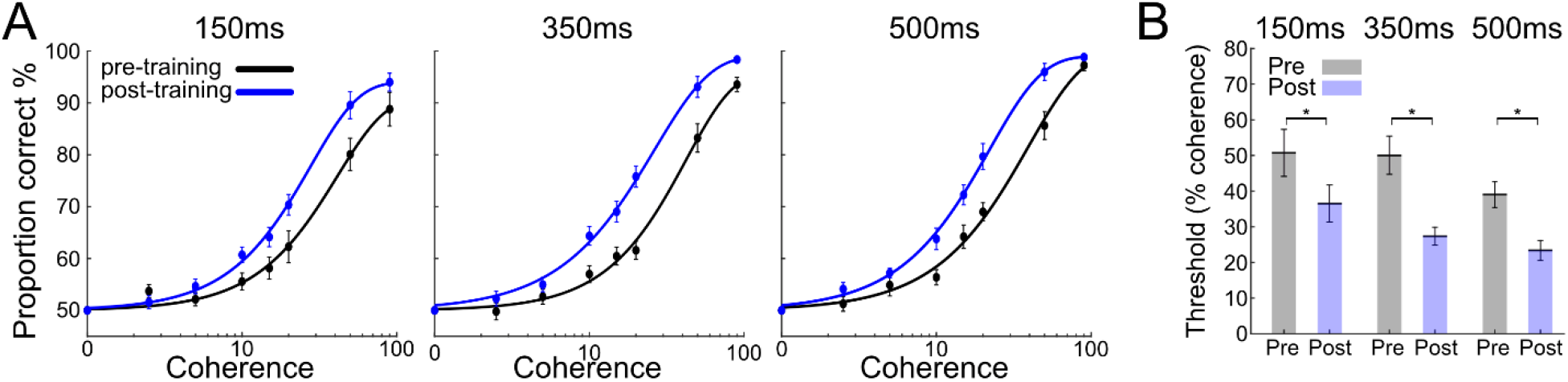
Training improved the performance of observers. A) Average psychometric functions of observers in each condition pre and post training in all three conditions. The observers’ performance as a function of coherence was characterized by fitting a Weibull function to the proportion of correct responses using the maximal likelihood algorithm. B) The mean psychophysical coherence threshold before and after the training in three different conditions. In all three conditions, observers’ thresholds were significantly reduced after the training. Threshold was defined as 82% correct of the Weibull function fits. Asterisks: statistically significant difference. Error bars show standard error of the mean (SEM).

To gain a more detailed understanding of how perceptual behavior changed in our observers, we analyzed two key aspects of their psychometric behavior: perceptual bias and the slope of the psychometric curve. To determine each observer’s perceptual biases, we first determined the proportion correct for both expansion and contraction stimuli. Having no bias would mean an equal tendency to choose expansion and contraction, with any deviation from this balance indicating a non-zero perceptual bias (see Methods for detail).

Figure 3A illustrates the biases of the observers pre- and post-training for each stimulus strength condition. In all conditions, observers showed a significant bias toward *contraction* before the training (150 ms (pre-training): mean of biases ± SEM = -0.261 ± 0.10; p = 0.0179; 350 ms (pre-training): mean of biases ± SEM = -0.183 ± 0.049, p = 0.011; 500 ms (pre-training): mean of biases ± SEM = -0.138 ± 0.077, p = 0.010, WSR test, Wilcoxon signed-rank (WSR) test; Fig. 3A). Post-training, these biases were unchanged in the 150 ms condition (mean of biases ± SEM = -0.147 ± 0.05; p = 0.0211), but were eliminated in the 350 ms condition (mean of biases ± SEM = -0.022 ± 0.045, p > 0.05). Interestingly, for the 500 ms condition, a significant bias toward expansion emerged with training (mean of biases ± SEM = 0.136 ± 0.040, p = 0.042, WSR test). Overall, in the 350 ms and 500 ms conditions, training resulted in a significant shift in bias (350 ms: a difference of biases = 0.264, p = 0.04; 500 ms, a difference of biases = 0.394, p < 0.01, WRS test) while in the 150 ms condition there was no significant change (150 ms: difference of biases = 0.125, p > 0.05, WRS test). Thus, training altered observers’ biases in different ways, depending on task difficulty.

**Figure 3.**
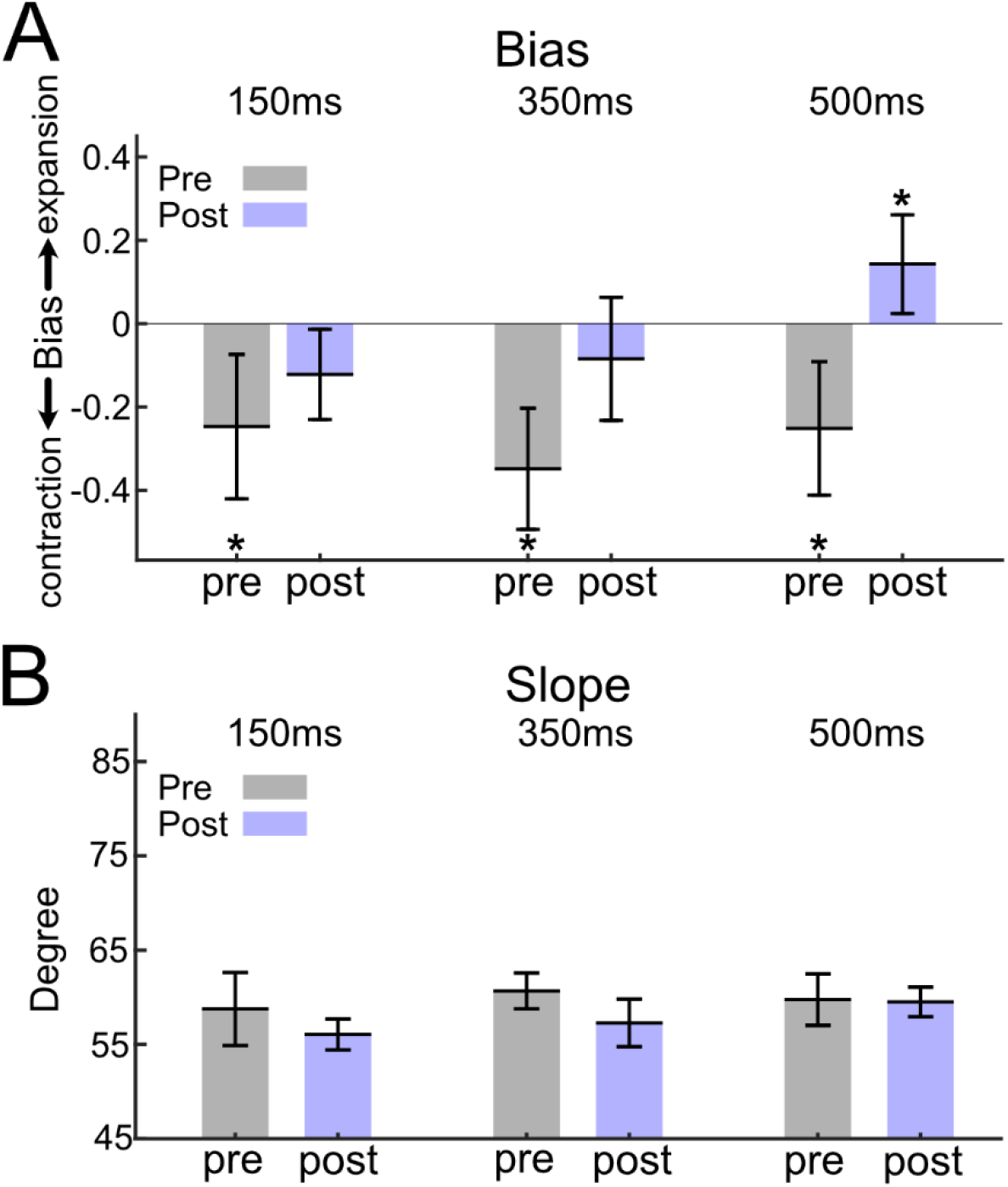
Changes in behavioral bias and slope of psychometric behavior before and after training. A) The average bias across observers in each condition, pre-training and post-training. Before training, the average bias in all conditions was significantly toward contraction. This contraction bias changed after training, with the average bias in the 500 ms condition significantly shifted toward an expansion bias. Negative bias values indicate a bias toward contraction, and positive values indicate a bias toward expansion B) The mean slope of psychometric behavior among observers did not change significantly pre and post-training across all conditions. Asterisks: statistically significant difference. Error bars show standard error of the mean (SEM).

We also compared psychometric curve slopes pre- and post-training. Any variation in the slope signifies a change in observer sensitivity to motion perception. However, on average, we observed no significant shift in slope for any of the three conditions (Figure 3B) (p > 0.05 for all three conditions, Wilcoxon rank sum (WRS) test).

The observed patterns of pre- and post-training biases suggest the use of a suboptimal readout and learning strategy, as biases were not always reduced through training. Indeed, for the easiest condition (500 ms), an expansion bias emerged with training, indicating that VPL did not optimize performance. An optimal readout and learning strategy, such as readout reweighting, would be expected to improve perceptual sensitivity while reducing biases to zero in all conditions, provided that observes received sufficient training (Petrov, Dosher, & Lu, 2005). In the following section, we employ computational modeling to evaluate various readout and learning algorithms based on their ability to replicate our experimental findings.

### Computational model

The psychophysical results presented in the previous section reveal the following pattern of perceptual effects (Figure 2): 1) In all conditions, observers showed strong contraction bias before training; 2) Training improved motion discrimination on average; and 3) Perceptual biases changed with training differently in the three conditions. In particular, in the most difficult condition (150 ms duration), the contraction bias remained after training. In the medium difficulty condition (350 ms), biases were typically reduced. In the easiest condition (500ms), a large expansion bias emerged with training.

To gain insight into how training improved performance and altered the perceptual biases of the human observers, we developed a neural network model that, similar to area MST in the visual cortex, had an asymmetrical sensory representation of the two stimulus conditions (contracting vs expanding optic flow). The two sensory populations (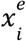 and 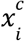) projected to a single readout neuron that outputs the decision variable (here, expansion or contraction decision; Figure 1A). The scalar output of the readout neuron was compared in the two stimulus conditions (i.e. expanding vs contracting optic flow), and the type of optic flow in each trial was determined based on a decision criterion (*c*_*d*_). If the decision neuron fired larger than *c*_*d*_, the input stimulus would be determined as an expanding stimulus and contracting otherwise. Since the majority of neurons in area MST are tuned to expansion optic flow, in our computational model, 80% percent of the sensory population were set to be selective for the expansion stimuli (Graziano, Andersen, & Snowden, 1994; Heuer & Britten, 2004). The tuning properties of the artificial neurons were fixed and did not change during training. The only trainable components of the model were the readout weights, which set the connection strength between the sensory neurons and the readout neuron, as well as the decision criterion (*c*_*d*_). The learning rules only modulated the readout weight and/or the decision criterion throughout training. The equation below summarizes the model:

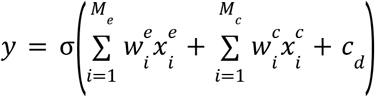

The goal was to find out which learning rule could best explain and reproduce observers’ behavior during the pre-training, training, and post-training phases. Before simulating the perceptual training experiment (our psychophysics experiments), we pre-trained the neural network model to simulate the pre-experiment condition of the human observers. To achieve this, we trained the model exclusively with high-strength optic flow stimuli, under the assumption that the human visual system is generally adapted to high-coherence optic flow encountered in daily life. Thus, the preliminary training phase was conducted using very high coherence level stimuli ranging from 0.5 to 1.0.

We investigated two learning rules during the initial training phase:

1. Optimal readout training: This learning rule adjusted the readout weights of individual sensory neurons (*w*_*i*_) as well as the decision criterion (*c*_*d*_) to achieve optimal performance.
2. Decision criterion modulation: This learning rule only adjusted the decision criterion while keeping the sensory readout weights unchanged.

Following the preliminary training, we evaluated the pre-trained model using stimuli with varying coherence levels (0.15 to 0.65), in order to mimic the pre-training test phase of our experiment. To simulate the three different stimulus durations used in the psychophysics experiments, we scaled the coherence levels by three distinct factors (0.3, 0.6, and 1). This scaling was done to incorporate the influence of duration in the model, as previously suggested (Law & Gold, 2008).

When we used the optimal learning strategy during the preliminary training phase, the model showed no perceptual bias when tested with low strength stimuli. This was inconsistent with the human observers’ pre-training contraction bias. In contrast, once the model was trained with the decision criterion modulation strategy, it consistently showed a significant contraction bias across all levels of difficulty. This outcome closely resembled the results obtained in the human experiment (Fig 4A; gray bars). These findings suggest that, with an asymmetric sensory population, a simple and suboptimal readout approach (i.e., modifying only the decision criterion) can effectively account for the initial contraction bias observed in our psychophysics observations.

**Figure 4.**
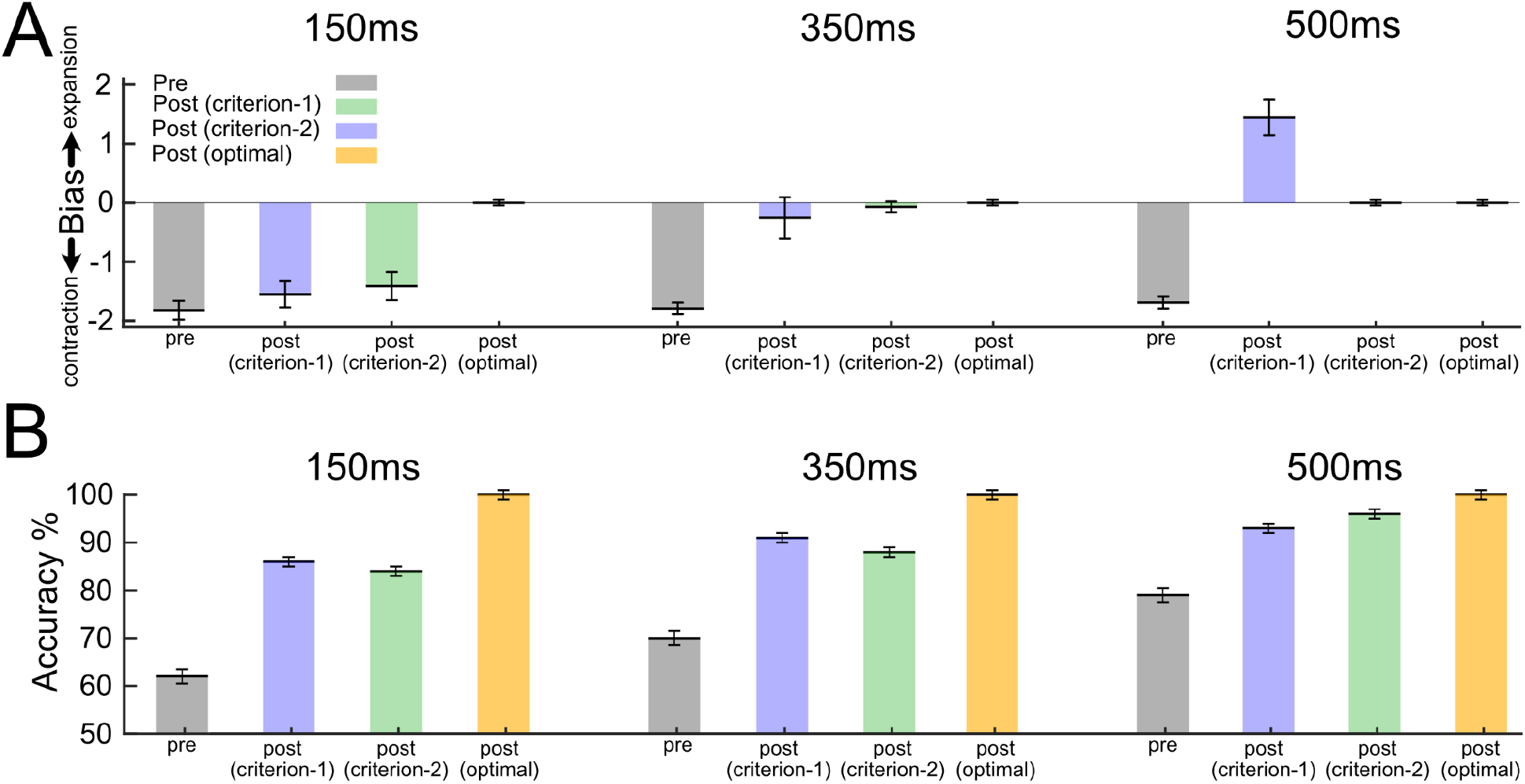
Comparison of the bias and discrimination accuracy of the model using different strategies during training. A) During the preliminary training phase, the model was trained with the decision criterion modulation strategy with high strength optic flow stimuli. The model showed a significant bias toward contraction, similar to the observers’ behavior (gray bars). After retraining with low strength stimuli, the decision criterion modulation strategy could reproduce human experimental results only when all sensory neurons in the population had the same readout weight (criterion-1; 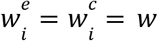) and not when the two populations had equal but opposite-signed readout weights (criterion-2; 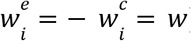). B) After the training, the discrimination accuracy (% correct) of all models were improved in all conditions. The accuracy of the optimal model always reached 100% after training. Error bars show the standard error of the mean (SEM).

Next, we simulated the nine-session training phase of our psychophysics experiments by re-training the model using low coherence stimuli ranging from 0.2 to 0.3. We specifically chose this low coherence range to mimic the 2-down 1-up staircase procedure employed during the nine-session training phase of the human experiment. This procedure required the observers to concentrate on a limited coherence range centered around their perceptual threshold. Again, note that the model can be trained for this phase either by using the optimal learning strategy or decision criterion modulation. Finally, similar to the post-training test in the human experiment, we tested the models again with the same range of coherences we used in the pre-training phase.

Following the training, all models exhibited improved discrimination accuracy across all conditions (Fig. 4B). However, the model employing the optimal learning strategy displayed no bias in any of the conditions after training (Fig. 4A). The decision criterion modulation, on the other hand, could fully reproduce the human biases: For the difficult condition, there was a slight reduction of contraction bias but the bias remained significantly toward contraction. For the moderate condition, the model had no significant bias after training. For the easy condition, significant bias toward expansion emerged after training. Notably, this expansion bias only occurred when all neurons in the population had the same readout weight 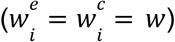;namely, the downstream readout neuron received a sum of their firing. An alternative approach, in which the two populations had equal but opposite-signed readout weights (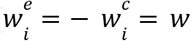 i.e. the downstream readout neuron receiving a difference of their firing), did not result in the same expansion bias.

To determine whether the asymmetrical sensory representation was necessary to account for these results, we tested the model using a symmetric sensory population. The decision criterion modulation was applied during the preliminary training, followed by decision criterion modulation or optimal learning strategies in the training phase. In this case, the symmetric sensory population was unable to reproduce human behavior (Fig. 5).

**Figure 5.**
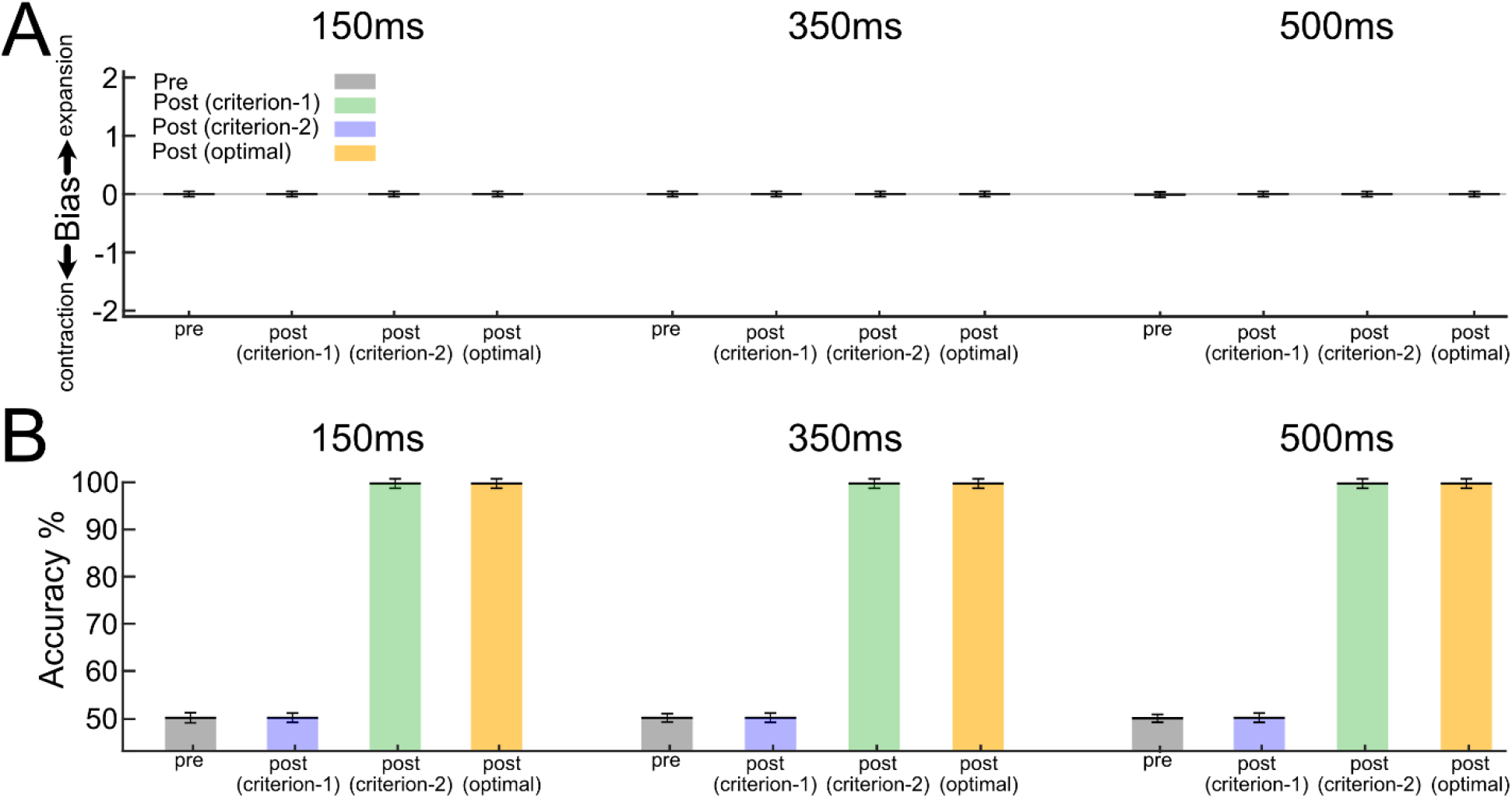
Comparison of the bias and discrimination accuracy of the model with a symmetrical population. A) We trained the model with a symmetrical population using the decision criterion modulation strategy during the preliminary training phase. In the pre-training test, the model did not develop a bias in any condition (gray bars). We then retrained the model with low strength stimuli using criterion-1 model (criterion-1; 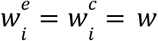; blue bars), criterion-2 model (criterion-2; 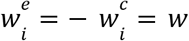; green bars) or optimal readout model (yellow bars). The bias of all models remained at zero in all conditions in the post-training phase. B) In the pre-training phase, the model’s discrimination accuracy (% correct) remained at the chance level. After the training (post-training phase), there was no improvement in the performance of the criterion-1 model while the discrimination accuracy of the criterion-2 and optimal readout models reached 100% in all conditions. Error bars show the standard error of the mean (SEM).

Humans can also enhance their performance by increasing their attention during the stimulus presentation. We can model attention in the cortex with an increased gain of sensory representations (Fox, Birman, & Gardner, 2023; Maunsell & Treue, 2006). Therefore, instead of altering the decision criterion, an alternative approach to training could involve adjusting the gain of sensory responses to align with the decision criterion value. We tested this strategy by introducing multiplicative gains to the sensory readout (*g*^*e*^ and *g*^*c*^ see Methods, equation 3). Instead of optimizing the readout weights or the decision criterion, we assessed the learning dynamics of adjusting these sensory gains during training. As shown in Fig. 6, the gain modulation strategy could reproduce our observed bias and accuracies across the three difficulty conditions. These findings indicate that modulating the decision criterion (which is also equivalent to an additive gain modulation of neurons’ responses; see Discussion for more details) and multiplicative gain modulation can reproduce our psychophysical results.

**Fig. 6.**
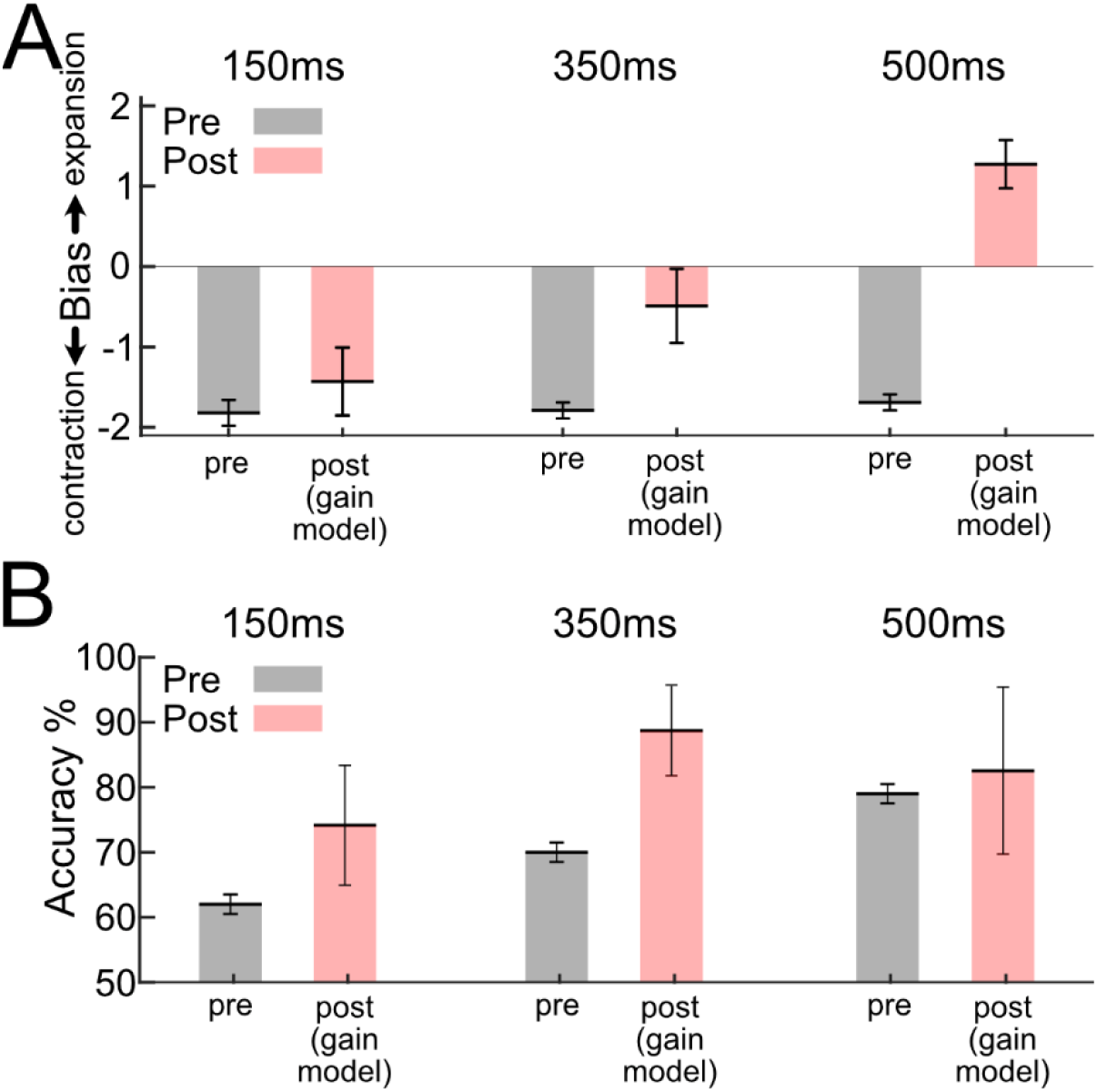
the bias and discrimination accuracy of the model using gain modulation strategy in the training phase. Perceptual training was modeled using the gain modulation strategy. The model could reproduce human behavior for (A) bias and (B) accuracy. Gray bars indicate the models’ results in pre-training, and red bars in post-training using the gain modulation strategy. Error bars show the standard error of the mean (SEM).

## Discussion

Practicing perceptual tasks has been shown to enhance performance, and these improvements have primarily been attributed to the adaptation of existing sensory representations or their readouts to the specific training task (Karni & Sagi, 1991; Lu & Dosher, 2022; Sotiropoulos, Seitz, & Series, 2011; Wenliang & Seitz, 2018). These adaptations are commonly assumed to be optimal, in the sense that the sensory and/or sensorimotor synaptic connections are expected to undergo changes that optimize task accuracy. However, in our study, we sought to challenge this *optimality* assumption by investigating perceptual learning within a paradigm that relied on an asymmetric sensory representation. The overrepresentation of specific classes of stimuli in visual cortex enables the possibility of a suboptimal readout solution, which is not viable with a symmetric sensory representation (Figure 1). For these stimuli, we find that observers exhibit patterns of biased perception that change with training (Figure 2) in a manner that depends on task difficulty (Figure 3).

Through computational modeling, we show that a simple adjustment of the decision criterion adequately accounted for the initial and the training induced biases of the observers (Figure 4). Notably, the alternative learning rules proposed in previous studies failed to explain the observed biases (Figures 5, 6). Considering the prevalence of asymmetric sensory representations in the cortex, our findings make critical contributions to our understanding of learning within the visual system. By uncovering the role of decision criterion modulation in perceptual learning, particularly in the context of asymmetric representations, we challenge the prevailing assumptions regarding optimal adaptation of sensory and sensorimotor connections.

### Representational asymmetries in visual cortex

The primate visual cortex consists of many different brain regions, most of which respond specifically to particular stimulus features. In most of these regions, the encoding of stimulus features is not uniform but rather biased toward specific stimulus properties. These biases can take the form of higher firing rates, narrower tuning curves, or non-uniform distributions of stimulus preferences. Asymmetries can also change across retinal locations (Albright, 1989; Ponce, Hartmann, & Livingstone, 2017; Sasaki, Rajimehr, Kim, Ekstrom, Vanduffel, & Tootell, 2006).

Probably the best-known example is the cardinal bias found for orientation selectivity in the primary visual cortex (V1) (DeValois, 1982; Li, Peterson, & Freeman, 2003): There are more V1 neurons that prefer horizontal or vertical orientations than oblique ones. This asymmetry might co-exist with a radial organization, in which the most common preferred orientation varies with receptive field position (Sasaki, Rajimehr, Kim, Ekstrom, Vanduffel, & Tootell, 2006). In either case, the result is that the V1 population response is greater for some orientations than for others, setting up the conditions for learning to exploit changes in decision criteria. This would seem to be important for the field of VPL, given that so many studies use oriented stimuli (Song, Sun, Wang, Zhang, Kang, Ma, Yang, Guan, & Ding, 2010).

Other biases that have been detected neurophysiologically include biases for dark stimuli (Komban, Kremkow, Jin, Wang, Lashgari, Li, Zaidi, & Alonso, 2014; Yeh, Xing, & Shapley, 2009), “daylight” colors (Lafer-Sousa, Liu, Lafer-Sousa, Wiest, & Conway, 2012), near disparities (DeAngelis & Uka, 2003; Tanabe, Doi, Umeda, & Fujita, 2005), small motion stimuli (L. D. Liu, Haefner, & Pack, 2016; Tsui & Pack, 2011), concave surfaces (Verhoef, Vogels, & Janssen, 2012), and curved shapes (Pasupathy & Connor, 2002). There are also entire brain regions dedicated to specific categories of stimuli, such as motion, color, faces, and food, and it is conceivable that observers rely on these asymmetries by changing their readout strategies during VPL (Bakhtiari, Awada, & Pack, 2020; Chen, Cai, Zhou, Thompson, & Fang, 2016; L. D. Liu & Pack, 2017).

These cortical asymmetries tend to reflect ecological or evolutionary experience (Brenner & Rauschecker, 1990; Mineault, Khawaja, Butts, & Pack, 2012). The strong preference for expansion over contraction optic flow is likely linked to the visual experience of frontal-eyed animals during locomotion, and the bias for small moving stimuli has long been associated with the function of separating objects from the background (Lettvin, Maturana, McCulloch, & Pitts, 1959; L. D. Liu, Haefner, & Pack, 2016; D. Tadin, Park, Dieter, Melnick, Lappin, & Blake, 2019). Cardinal orientations are apparently overrepresented in natural images (Girshick, Landy, & Simoncelli, 2011), and the bias toward curved or circular stimuli in mid-level cortex could be part of a network for face processing in higher-level cortex (Ponce, Hartmann, & Livingstone, 2017). Our findings suggest that the brain readily leverages these asymmetries during VPL, highlighting the importance of neurophysiological considerations in visual learning (Bakhtiari, Awada, & Pack, 2020).

In this regard, it is worth considering the influence of natural stimuli outside the laboratory during learning experiments. For experiments that span several days, exposure to the natural world would be expected to counteract or reinforce VPL, depending on the nature of the stimuli and their representation in cortex.

### Comparison with previous psychophysical results

Some of the abovementioned cortical asymmetries have likely perceptual consequences. For orientation, there is the oblique effect (Appelle, 1972), in which human and animal observers are more sensitive to horizontal and vertical orientations than to others. As with cortical responses, human psychophysical observers are more sensitive to dark stimuli than to light stimuli (Komban, Kremkow, Jin, Wang, Lashgari, Li, Zaidi, & Alonso, 2014). The cortical preference for small moving stimuli is associated with a wide range of perceptual phenomena presumably related to figure-ground segregation (D. J. V. r. Tadin, 2015). The overrepresentation of foveal stimuli in most of retinotopic cortex has been linked to perceptual effects that occur during eye movements (Richard, Churan, Guitton, & Pack, 2009). And the overrepresentation of entire classes of stimuli, such as faces, is thought to be partly responsible for illusory perceptual phenomena (i.e., pareidolias). Some of these biases can be reduced or abolished with training (Bakhtiari, Awada, & Pack, 2020; Furmanski, Schluppeck, & Engel, 2004).

An example of this phenomenon was recently presented in (Szpiro, Burlingham, Simoncelli, & Carrasco, 2022), where it was demonstrated that perceptual learning of orientation discrimination amplified the pre-existing bias towards cardinal orientations. They modeled the post-training improvement in perception by proposing an increased gain of neurons encoding the task stimuli which could also reproduce their observed humans’ biases. However, an important distinction between their study and ours lies in the fact that their task stimuli consisted exclusively of under-represented stimuli, which could lead to different perceptual biases.

The consequences of these asymmetries are seldom considered in VPL studies, most of which are concerned with changes in sensitivity. Indeed, many models assume that decision criteria are set optimally, in an unbiased manner (Petrov, Dosher, & Lu, 2005). Nevertheless, in detection paradigms, people often alter their decision criteria, even though this is not the optimal strategy (Wenger, Copeland, Bittner, Thomas, & Psychophysics, 2008), and this can result in biased perceptual responses (Seitz, Nanez, Holloway, Koyama, & Watanabe, 2005). Our findings extended this strategy to a discrimination task that involved asymmetrical cortical representations of two distinct stimulus conditions. In this particular case, humans appear to be able to transform the discrimination task into a detection task, by aiming to detect the overrepresented stimulus condition (here, the expansion optic flow).

Some previous work has examined the effects of asymmetries in the design of perceptual learning tasks. In these experiments, asymmetries are created by presenting one stimulus more often than another, or by providing false or irrelevant feedback (Herzog, Ewald, Hermens, & Fahle, 2006; Herzog & Fahle, 1999; Seitz, Nanez, Holloway, Koyama, & Watanabe, 2005). In these cases, observers appear to respond by altering their decision criteria. While changes in sensitivity and in criterion often occur simultaneously (Wenger, Copeland, Bittner, Thomas, & Psychophysics, 2008), they have different characteristics, the most notable of which is the faster dynamics of changes in criteria (Aberg & Herzog, 2012).

### Implications for learning in the cortex

In most psychophysics experiments, it is challenging to differentiate between different strategies and learning rules. A change in decision criterion would likely yield results similar to improved attentional focus, which is known to be crucial for learning (Szpiro & Carrasco, 2015). Specifically, a modulatory attentional effect that changes the excitability of sensory neurons’ responses could be similar to the decision criterion modulation observed in our study. Interestingly, previous research has reported biases in perceptual responses to attended stimuli versus unattended stimuli (Itthipuripat, Chang, Bong, & Serences, 2019) showing a perceptual effect similar to the observations in this study.

Despite the challenges of distinguishing between different learning strategies, our results are in conflict with the prevailing notion that performance is optimized through learning. This notion may hold true when using stimuli with symmetric representations and investigating learning solely based on perceptual sensitivity (Petrov, Dosher, & Lu, 2005). However, under asymmetric stimulus representations, which are common in the cortex, as mentioned earlier, suboptimal and easier solutions may be adopted (J. Liu, Dosher, & Lu, 2015). Our study shows that an optimal learning algorithm, such as gradient descent, does not automatically discover the suboptimal and easier solution under asymmetric representations.

Fitzgerald et al. (2013) raised an intriguing possibility that observers may prefer to create low-dimensional representation of the task stimuli. This enables observers to introduce their own representational asymmetries, even where the underlying sensory representations of the task stimuli are symmetric. The creation of such low-dimensional and asymmetric representations, as demonstrated in this study, facilitates a simpler learning process for the stimulus-response mapping. Therefore, learning strategies that exploit asymmetric representations may potentially remain latent in numerous VPL experiments, including those that do not explicitly rely on the intrinsic representational asymmetries of the visual cortex.

For certain kinds of learning tasks, asymmetric representation could be useful. For applications of VPL in particular, changes in sensitivity often require prolonged training and fail to generalize beyond the trained stimulus, limiting their utility for therapeutic purposes or other applications. Our results, along with previous findings (Aberg & Herzog, 2012) suggest that decision criteria can be adjusted more rapidly, often within a single session, although the learning effects might also be more transient (Aberg & Herzog, 2012). One goal for future research could therefore be to exploit representational asymmetries in the brain to develop faster learning (Pandey, Neupane, Vaidya, Adhikary, & Pack, 2022).

### Comparison with previous computational models

Previously proposed theories of VPL have primarily suggested *optimal* changes either in the sensory representations (referred to as re-tuning; (Karni & Sagi, 1991; Wenliang & Seitz, 2018)) or in the readout of sensory neurons (referred to as re-weighting; (Lu & Dosher, 2022; Sotiropoulos, Seitz, & Series, 2011)). However, these theories have assumed homogeneous sensory representation of task stimuli, while, as we showed in this paper, an asymmetric representation enables a simpler learning strategy that does not align with the existing literature. Recent studies have provided computational evidence demonstrating that artificial neural networks (ANNs) also display similar representational asymmetries, as they mirror the biases present in their training data (Benjamin, Zhang, Qiu, Stocker, & Kording, 2022). Learning algorithms capable of leveraging these representational asymmetries may bring ANNs closer to capturing human learning dynamics.

For a naïve observer, before training, the readout weights of sensory neurons are not inherently different due to the unfamiliarity with the task and the absence of a priori assumptions regarding the relative importance of these neurons in the task (Wenliang & Seitz, 2018). In the case of asymmetric representation of task stimuli, as discussed earlier, the initial equal readout weights are effective enough to solve the task (Figure 1). The key question is whether humans would optimize learning by adjusting the readout weights of all sensory neurons, deviating from equal readout weights, or if they would maintain the suboptimal strategy of equal readout weights and enhance performance solely by modulating the decision criterion. Our findings suggest that, when feasible, humans learn to improve task performance by solely adjusting the decision criterion, without modifying the readout weights. It is also justifiable from a computational complexity perspective as it is more efficient to find and store the optimal value of one parameter (i.e., the decision criterion) rather than optimizing a large number of sensory readout weights. This becomes more critical for multi-task learning. Previous work on few-shot learning in ANNs (Triantafillou, Larochelle, Zemel, & Dumoulin, 2021) has suggested the advantages of a similar approach in machine learning; namely, keeping the feedforward connection weights stable and adjusting the bias and scale of the readout from each layer for every new task. Although our focus in this study was primarily on sensorimotor readout, this approach could potentially be applicable to multiple layers.

Furthermore, our observed modulation of the decision criterion can be interpreted as a global change in the firing rate of sensory neurons. Specifically, a decrease (or increase) in the decision criterion in our model corresponds to a global increase (or decrease) in the firing rate of sensory neurons (see equation 3). We also showed that learning via adjusting a multiplicative gain could reproduce the biases observed in humans. These results suggest that the proposed role of decision criterion or gain modulation in perceptual learning could be implemented through an additive or multiplicative top-down modulation of the global firing rate of sensory neurons. Interestingly, a previous neural network model of perceptual learning (Herzog & Fahle, 1998) highlighted the significance of top-down modulation signals in capturing the dynamics of perceptual learning. This observation implies that, unlike gradient descent, more biologically plausible learning algorithms that rely on top-down modulation of neurons’ activation, such as target propagation (Lee, Zhang, Fischer, & Bengio, 2015) and its more recent variants (Meulemans, Zucchet, Kobayashi, von Oswald, & Sacramento, 2022), may be able to reproduce our observations. Despite converging to similar solutions in the steady-state (Meulemans, Zucchet, Kobayashi, von Oswald, & Sacramento, 2022), the trajectory of reaching the final solution may differ among these learning algorithms which make comparisons with the dynamics of human learning worthwhile.

In our modeling experiments, we trained all variants of the model for a fixed number of epochs, considering the time constraints in human psychophysics experiments. After this fixed training period, we compared the performance of different learning strategies. However, we also noticed that longer training times for two of our learning strategies, namely decision criterion modulation and gain modulation, resulted in distinct bias patterns. Unlike decision criterion modulation, extending the training period with gain modulation led to an expansion bias across all task conditions (easy, medium, difficult). Currently, our psychophysics data are insufficient to differentiate between these two possibilities, as longer training periods are necessary for the more challenging conditions.

## Conclusion

We demonstrated that, when relying on asymmetric sensory representations, humans use a simple readout and learning strategy to improve their perceptual performance. This strategy cannot be explained by the optimal learning algorithms proposed in previous studies.

Considering the widespread representational asymmetries in the cortex, our findings carry significant implications for learning mechanisms in cortical processing. Future research should explore a broader range of learning algorithms, particularly those that incorporate top-down feedback modulations, to examine their capacity to replicate the learning dynamics and biases observed in humans.

## Acknowledgements

This work was funded by NSERC Discovery grants to CCP (RGPIN/04333-2017) and SB (RGPIN-2023-03875).

